# Highly Adaptive Conductive Polymer Electronics Enhance Neural Data and Learning Accuracy

**DOI:** 10.1101/2025.10.14.682347

**Authors:** Vittorio Mottini, Liuxi Xing, Charlie Meilinger, Soham Inamdar, Xiang Calvin Chen, Jiaqi Wang, Yi Xing, Jack Darbonne, Zhengxu Tang, Yunnuo Zhang, Christopher H. Contag, Mi Zhang, Jinxing Li

## Abstract

Human skin, the body’s largest organ, plays a vital role in sensing and transmitting neuronal, mechanical, and biochemical signals, making it an essential non-invasive interface for health monitoring, rehabilitation, and human-machine interaction. However, aging-related changes, including thinning, increased wrinkling, dryness, and altered collagen structure, significantly impact electrical impedance, conductance, and contact stability, challenging the fidelity and consistency of bioelectronic signal acquisition. Here, we address this gap by developing “AdapSkin,” an age-adaptive, skin-mimicking, bio-adhesive, and stretchable polymeric electronic skin interface that seamlessly conforms to diverse skin properties, enabling high-fidelity and high-density electrophysiological recording. The soft electrodes of AdapSkin are composed of an aqueously processed, homogeneously mixed organic nanocomposite with a conductive polymer percolation network, forming a gel-like interface that reduces modulus and enhances skin-electrode contact. The materials platform achieves extraordinary softness and electrical stretchability of up to 1200% through a double-network composite structure. AdapSkin significantly minimizes age-induced variations in interfacial impedance and signal-to-noise ratio (SNR), improving signal consistency for neuromuscular assessment, prosthetic control, and rehabilitation applications. Scalable fabrication enables the creation of large-area electrode arrays, which reduces motion artifacts, improves sEMG mapping reliability, and ensures long-term signal stability across various age groups. Machine learning analysis further demonstrates AdapSkin’s superior accuracy in gesture classification for elderly users, highlighting its potential to enhance prosthetic control, assistive robotics, and rehabilitation for individuals with sarcopenia and neuromuscular decline. By improving signal quality and adaptability in aging populations, AdapSkin advances fair bioelectronic interfaces, fostering more equitable and effective healthcare technologies for age-related conditions.

## Main Text

Beyond its primary role as a protective barrier, human skin connects inner physiology and the external environment, and can therefore play a multifaceted role in monitoring health and facilitating human-machine interactions (*1*–*6*). Skin-interfaced bioelectronic devices (electronic skin or e-skin) can non-invasively monitor the neuronal and mechanical signals from muscles and various organs through the epidermis (*1*–*5, 7, 8*), enabling the diagnosis of various muscular, cardiac, or neurological conditions, and the control of prosthetic devices or robots in real-life or virtual reality (VR) environments (*9*–*13*). Recent advances in flexible electronics and sensing technologies have also led to skin-like and skin-interfaced biomolecular sensors for continuous, non-invasive screening of metabolic analytes through sweat (*14*–*17*). This emerging skin-interfaced bioelectronics shows incredible potential to enhance health monitoring, early disease prediction, and human augmentation, leading to scalable and ubiquitous healthcare for precision medicine (*5, 18*–*20*).

However, the surface properties of skin vary individually, and age over time. For example, compared to young individuals, the skin of senior people becomes thinner with noticeable wrinkles and age spots associated with reduced strength and elasticity (*21*). Their sebaceous glands produce less oil, while the sweat glands also produce less sweat, making it harder to keep the skin moist, resulting in dryness and itchiness (*22*–*24*). These biomechanical and compositional differences alter skin electrical impedance and conductance (*25*–*28*), as well as its contact with electronics, and hence challenge precise and consistent signal recording and analysis, as well as the adaptability and fairness of current skin-interfaced bioelectronics. For example, surface electromyography (sEMG), produced by muscle fiber firing in response to neural signals, is widely used to evaluate muscle function, diagnose neuromuscular disorders, monitor rehabilitation techniques, and serve as a non-invasive neural interface for augmented reality (AR) and VR environments (*29, 30*). Non-invasive sEMG recordings are particularly desired for elderly individuals with sensitive or fragile skin for biofeedback to improve the efficacy of rehabilitation and disease management (*31, 32*). Nevertheless, current sEMG interfaces, using metallic dry electrodes or electrolytic gel electrodes that can dry out easily overtime, usually cause the rigid electrodes to lose conformal contact with wrinkled and uneven skin surfaces during use and motion (*33*). These issues are more significant for seniors with wrinkled or dry skin, leading to increased interface impedance, motion artifacts, signal drift, source dislocation, and skin irritation over time (*34*–*36*). Due to the aging of motor neurons (*37*), atrophy and loss of muscle mass (sarcopenia) (*38*), improved bioelectronic interfaces are crucial for seniors to enable high-fidelity, high-density, high-stability, and high-comfort biodata acquisition for precision health restoration and maintenance (*39*–*41*).

To improve the fairness of wearable technologies and skin-interfaced electronics across the population, it is essential to create interfaces that can conform to the skin’s features without discrimination and adapt to the user during routine activities. Building upon recent advancements in soft and stretchable bioelectronics (*1*–*5*), we address this technical gap by developing “AdapSkin,” an intrinsically soft and bio-adhesive sensor array that offers seamless interfacing with varying skin characteristics for high-fidelity surface electrophysiological signal recording (Fig. 1A). The high stretchability and conductivity of the sensing materials is achieved a double-network composite structure, formed by mixing the conductive polymer poly(3,4-ethylenedioxythiophene) polystyrene sulfonate (PEDOT:PSS) polymer and the highly elastic polyurethane. Glycerol, a biocompatible material widely used in daily skin care (*42*), is added to enhance the stretchability, reduce impedance, and promote bio-adhesion. The In-Skin electrode array is highly stretchable and conforms seamlessly to the complex three-dimensional (3D) skin geometry during muscle contractions and relaxations due to the conductor’s high softness and adhesion. sEMG recording across a wide range of participants shows that AdapSkin significantly reduces the variation of data quality across populations with different skin characteristics, such as wrinkles and hairs, with low motion artifacts and stable continuous recording over time. Machine learning for gesture classification based on sEMG data indicates that AdapSkin achieves significantly higher accuracy in gesture recognition than current sEMG electrodes, particularly for elderly individuals with wrinkled skin. Our work highlights the challenges of effective rehabilitation therapies for senior individuals, whose EMG signals are not only weakened by the degradation of motor neurons but also compromised by skin degradation. Our work suggests that high-fidelity interface design will not only enhance the fairness and adaptability of electronics but also significantly contribute to the intelligence and accuracy of future machine learning models. Overall, our study addresses a critical gap in existing sEMG technology by offering improved signal quality and adaptability, thereby expanding the inclusivity of human-machine interfaces.

**Fig. 1.**
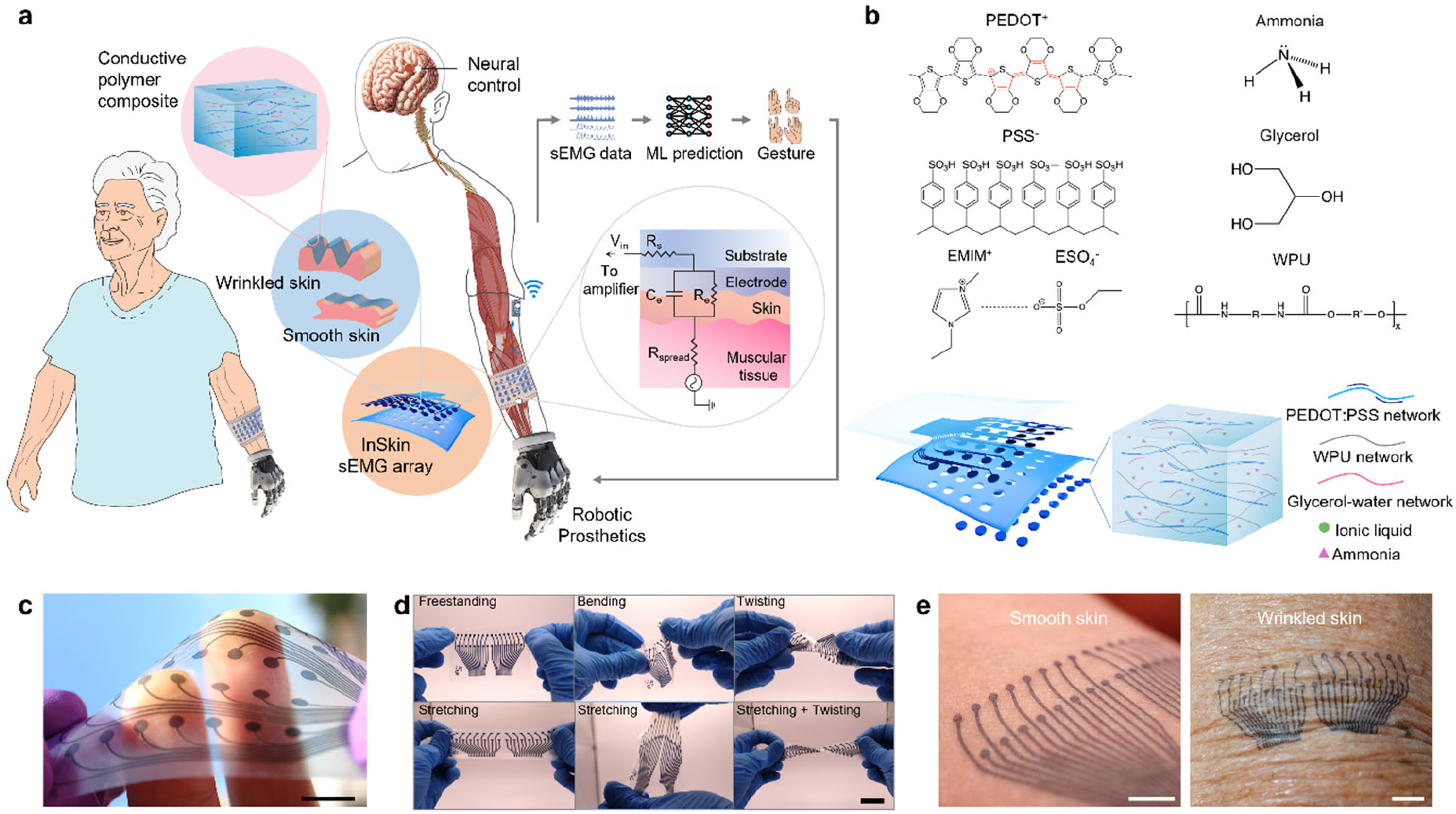
AdapSkin: an age-adaptive human machine interface made through intrinsically stretchable polymeric electronic skin. (**A**) Overview of the adaptive myoelectric interface system, including illustrations of the stretchable conductor design, the skin-electrode interface, and the multilayer device, showing how AdapSkin stretchable interface can conform to diverse skin topographies for high-fidelity EMG data recording towards precision gesture recognition and robotic control. (**B**) Molecular structure of the chemicals used for making the polymeric stretchable conductor, and illustration of the multichannel electrode array and its compositions. (**C**) Photograph of a 32-channel (4×8) AdapSkin stretchable electrode array. (**D**) Photographs a 32-channel (2×16) AdapSkin stretchable electrode array showing the deformability under twisting, stretching, and complex deformation. (**E**) Photograph of an array of the stretchable conductor laminated on the skin with different levels of wrinkles, highlighting the conformality of the stretchable composite with the skin’s complex surface features, scale bar: 5mm.

### Materials Design and Device Fabrication

To enable adaptability to different skin types, a solution-processable, conductive polymer composite is engineered to create the AdapSkin electrode array, which features high mechanical softness, low electrical impedance, and bio-adhesion. Such material design is expected to form a conformal interface and durable contact on uneven, moving, and deforming skin surfaces. The key material consists of a polymer mixture of PEDOT: PSS with water-borne polyurethane (WPU) to form a conductive polymer percolation network with excellent conductivity and stretchability (Fig. 1B). Such a method by blending the PEDOT:PSS dispersion with WPU offers an efficient approach in creating elastic and conductive polymer composites while maintaining the advantages of liquid processing for manufacturing large-area electronics (*43*–*45*). Ionic liquids (IL), 1-ethyl-3-methylimidazolium cation, and ethyl sulfate anion (EMIM:ESO_4_), were added to serve as non-volatile plasticizers, while a small amount of ammonia was added to prevent acid-induced aggregation of the anionic WPU by protonation from PSSH of the PEDOT: PSS (*45*–*47*). Particularly, we identified that glycerol, a widely used chemical in skin care and wound treatments, can significantly enhance the electrical stretchability (conductivity *vs* strain) and bio-adhesiveness of the polymer matrix (see details in the subsequent discussion and Methods section)

The substrate and encapsulation layers can be made of a commercially available, semi-permeable, self-adhesive, and elastic polyurethane film with an acrylic adhesive (Tegaderm™, thickness 27 μm), which is widely used as a biocompatible dressing to protect wounds and catheter sites. A laser patterning and transferring process is developed to fabricate the epidermal sEMG biopotential sensor array, comprising a multilayered assembly of intrinsically stretchable materials. The stretchable conductor is processed as a thin film through spin-coating, patterned as a multielectrode array using CO_2_ laser engraving, and then transferred and sandwiched between two polyurethane layers (fig. S1). This process allows scalable fabrication of electrode arrays with a device size of up to 12 inches (with the smallest feature size of approximately 50 μm), overcoming the size limitations of conventional photolithography processes with a potential for whole arm or even whole-body EMG mapping (fig. S2). Directly spin-coating polyurethane polymer could further reduce the thickness of the elastic substrate and the encapsulation layer. The active components made with the polymer composite after water evaporation can be as thin as 15 µm, while the substrate and encapsulation layer can each be 7.5 µm thin using spin-coated polyurethane film, making the device thickness lower than 35 µm (fig. S3). Figs. 1C-D show the AdapSkin array and its high mechanical compliance in bending, stretching, and twisting. Particularly, the conductive polymer conforms to complex skin microscale morphologies, such as different levels of wrinkles (Fig. 1E), and can be patterned to fit various body locations (fig. S4). The electrode’s softness and thinness allow seamless conformity to underlying microscopic skin features and skin hairs, as evidenced in the SEM images in fig. S5.

### Polymer Electric Materials Characterization

We first optimized the electromechanical properties of the stretchable polymer conductor, and it was noticed that glycerol, widely used in skin care for moisture, plays a critical role in the homogeneity and stretchability of the materials. The glycerol with multiple hydroxyls (-OH) groups can form glycerol-water networks through hydrogen bonding with water (*48*), which weakens the polymer chain interaction of the PEDOT:PSS, thus promotes the polymer dispersion in the polymer mixture and prevents their aggregation and consequent composite embrittlement (Fig. 2A). Maintaining water content in the polymer network also leads to the gelation of the materials and further increases the softness of the materials and the adhesion to the skin. To optimize the glycerol concentration, a water-borne conductive polymer dispersion mixture (referred to as solution CP, consisted of PEDOT:PSS, ionic liquid, waterborne polyurethane, and ammonia mixed in a volume ratio of 124:186:23:1, see details in Supplementary Materials) was optimized first based on their solubility and conductivity (249 S/m, fig. S6, see details in Supplementary Materials), and polymer solutions with different CP-glycerol weight ratios, ranging from 1:1 to 1:300, are then prepared to make thin films using spin-coating. Under 100% strain, cracks perpendicular to the stretching direction are observed for the film made without glycerol due to the PEDOT:PSS aggregation, while adding glycerol (CP-G 250, CP-G refers to the mixture of conductive polymer with glycerol, while 250 indicates the CP-glycerol volume ratio is 250:1, and so forth) significantly reduces cracks formation and leads to a homogenous smooth film under stretching (Figs. 2B). SEM images show obvious phase separation between the PEDOT:PSS and polyurethane while adding glycerol leads to a homogeneous polymer film (Fig. 2C). The glycerol-water networks also reduce the modulus of the materials and make the materials more stretchable, with freestanding films exhibiting no fracture up to 500% elongation (Fig. 2D). The electromechanical stretchability of the composite on the polyurethane substrate was characterized by evaluating the relative change in resistance (*R/R*_*0*_) under tensile stretching. All materials exhibit an increase in resistance, with CP-G 250 showing the lowest increase when stretched with a 500% strain and can be stretched under a strain of 1200% with an *R/R*_*0*_ smaller than 50 (Figs. 2E-F). Therefore, CP-G 250 is used to fabricate the AdapSkin electrodes. CP-G 250 also shows excellent electrical stretchability during cycling with a 100% strain (Fig. 2G). Interestingly, the composite could withstand 100 stretching cycles at 100% with an initial *R/R*_*0*_ of 1.47 and a final ratio of 0.93, showing a strain-insensitive property. Cyclic tensile strain characterization shows hysteresis loops characteristic of viscoelastic materials for both materials with and without glycerol, and the materials show a very low modulus smaller than 500 kPa (Figs. 2H). Once the stretchable conductor is transferred onto the polyurethane elastomer substrate, then the mechanical properties of the stacked materials are dominated by the elastomer, as demonstrated by the lower hysteresis (Figs. 2I). The electrode impedance of the materials was characterized and compared with commercial electrodes (CEs, 3M, FDK001commercial Ag/AgCl coated with sticky gel, sensing area 3.48 cm^2^) of the same size (Figs. 2J-L). When tested in a PBS buffer (pH 7.4), the polymer composite electrodes showed a significant reduction in impedance of 93.8%. The reduced impedance of the polymeric electrode is due to (1) the dual conductive nature (both electronically and ionically conductive) of PEDOT:PSS, (2) the nanoporous structure of PEDOT:PSS enabling high volumetric capacitance, and (3) the interconnect volume of the glycerol-water networks and conductive polymer networks further reducing the interfacial impedance without changing the electrode surface area as the electrolyte diffuses into the polymeric interconnect (*49, 50*). Comparing the average absolute impedance values within the frequency interval of interest for sEMG signals (20-500Hz), polymer CP and CP-G 250 electrodes showed 70% and 89% lower average impedance than CEs (Fig. 2L). Cyclic voltammetry characterization showed that having glycerol in the polymer improves the volumetric capacitance of the electrode by 132.7% Fig. 2M (fig. S6), and the impedance increases only 50% under a 100% strain (Fig. 2N).

**Fig. 2.**
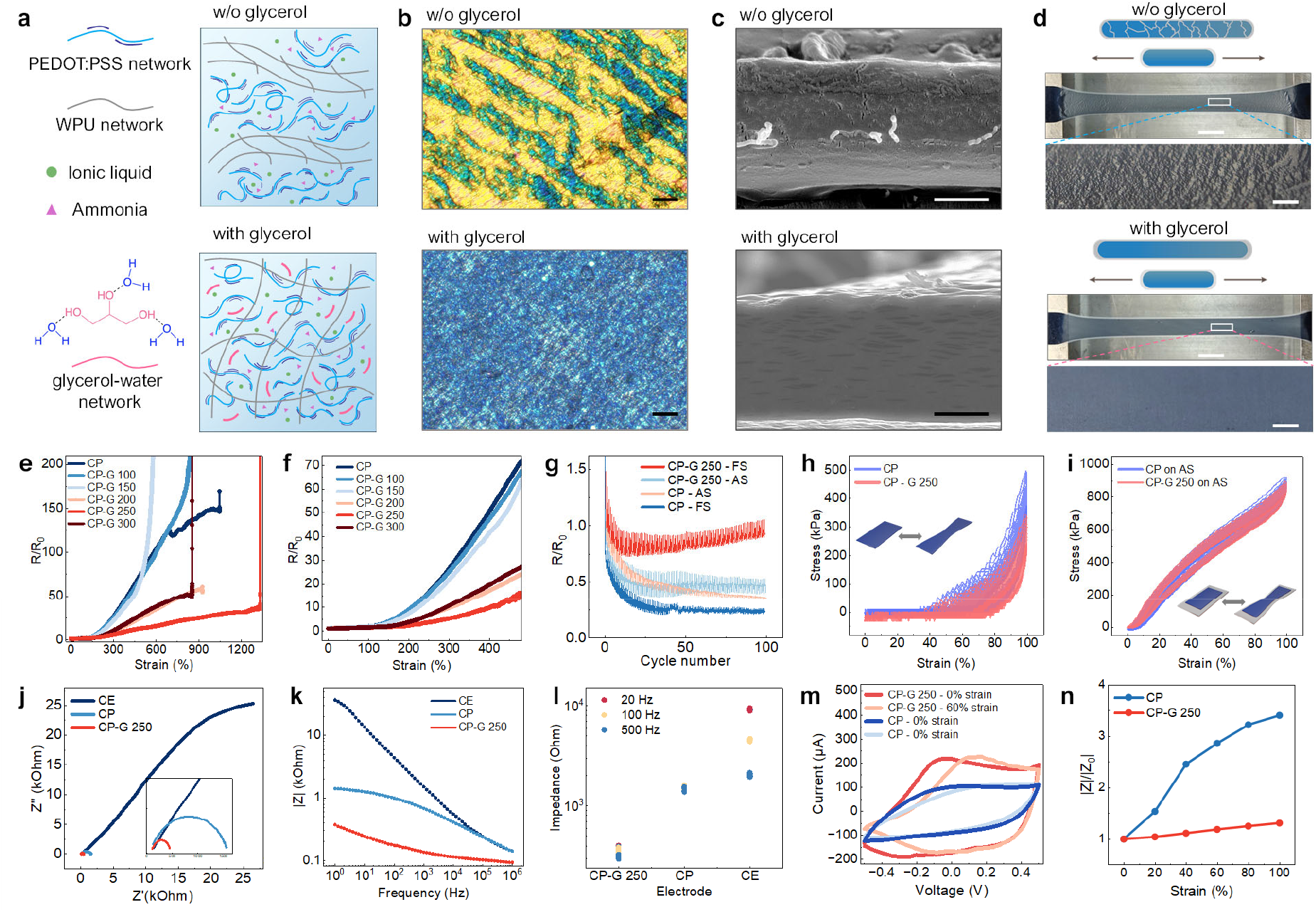
Morphological and electromechanical characterization of the intrinsically stretchable electronic skin polymer films. (**A**) Illustrations of the conductive polymer network with and without adding glycerol. (**B**) Polarized microscopy images of the conductive polymer film CP (top) and WP-G 250 (bottom) under a 200% uniaxial stretching, Scale bars: 100 µm. (**C**) Scanning electron microscopy images showing the cross-section of the CP and CP-G 250 polymer thin films. Scale bars: 5 µm. (**D**) Photos and illustrations of samples in polymer CP (top, without glycerol) and polymer CP-G 250 (bottom, with a CP-glycerol volume ratio of 250:1) under a 500% uniaxial stretching, showing the formation and absence of cracks, respectively. Scale bars: 10 mm (upper); 1 mm (lower). (**E-F**) Normalized strain induced resistance change for the polymer CP and Solution CP-G 250. (**G**) Cyclic strain-induced resistances change for free-standing CP, CP-G 250, and polyurethane-supported CP, CP-G 250. (**H-I**). Cyclic stress-strain curves for free-standing CP and free-standing CP-G 250 (H), and polyurethane-supported CP and polyurethane-supported CP-G 250 (I). (**J-K**) Electrochemical impedance spectroscopy Nyquist plot (J) and Bode (K) of Ag/AgCl commercial electrode, and electrodes made by CP and CP-G 250. (**L**) Electrode impedance for selected frequencies of relevant interest for sEMG signal detection (20 Hz, 100 Hz, and 500 Hz) of Ag/AgCl commercial electrode (CE) and electrodes made by CP and CP-G 250. (**M**) Cyclic voltammograms recorded in PBS buffer (pH 7.4, scan rate 50 mV/s) during a uniaxial stretch cycle to 60% strain (more details in fig. S6). (**N**) The relative impedance of the strained soft electrodes made by CP and CP-G 250 at 250 Hz.

### Adaptive Skin Biointerfacing

We then evaluated the AdapSkin electrode array for high-density sEMG recording and mapping. Compared with commercial electrode arrays, which have low resolutions, the 32-channel (4×8) AdapSkin sEMG arrays, featuring closely spaced electrodes, allow for the simultaneous recording of EMG signals from multiple muscles, enabling accurate and reliable non-invasive high-resolution sEMG mapping (Fig. 3A and Supplementary Movie 1). The fabricated 32-channel AdapSkin array has a narrow impedance amplitude distribution, indicating the high reliability of the fabrication method (fig. S7). The electrode array (2×16) can also easily identify muscle innervation zones (Fig. 3B), where the nerve and muscle connect. These capabilities provide critical insights into muscle function and coordination, enabling a comprehensive understanding of the underlying neuromuscular dynamics and contractile patterns, with implications for studying age-related muscle changes and providing precise biofeedback for rehabilitation. The AdapSkin electrodes can also be used to measure high-quality electroencephalogram (EEG) and electrocardiogram (ECG) (fig. S8), and remain securely attached to the skin for up to 24 hours while recording sEMG signals consistently, with only a 25% loss of SNR, indicating their potential for diverse applications in clinics and human-machine interactions (fig. S9).

**Fig. 3.**
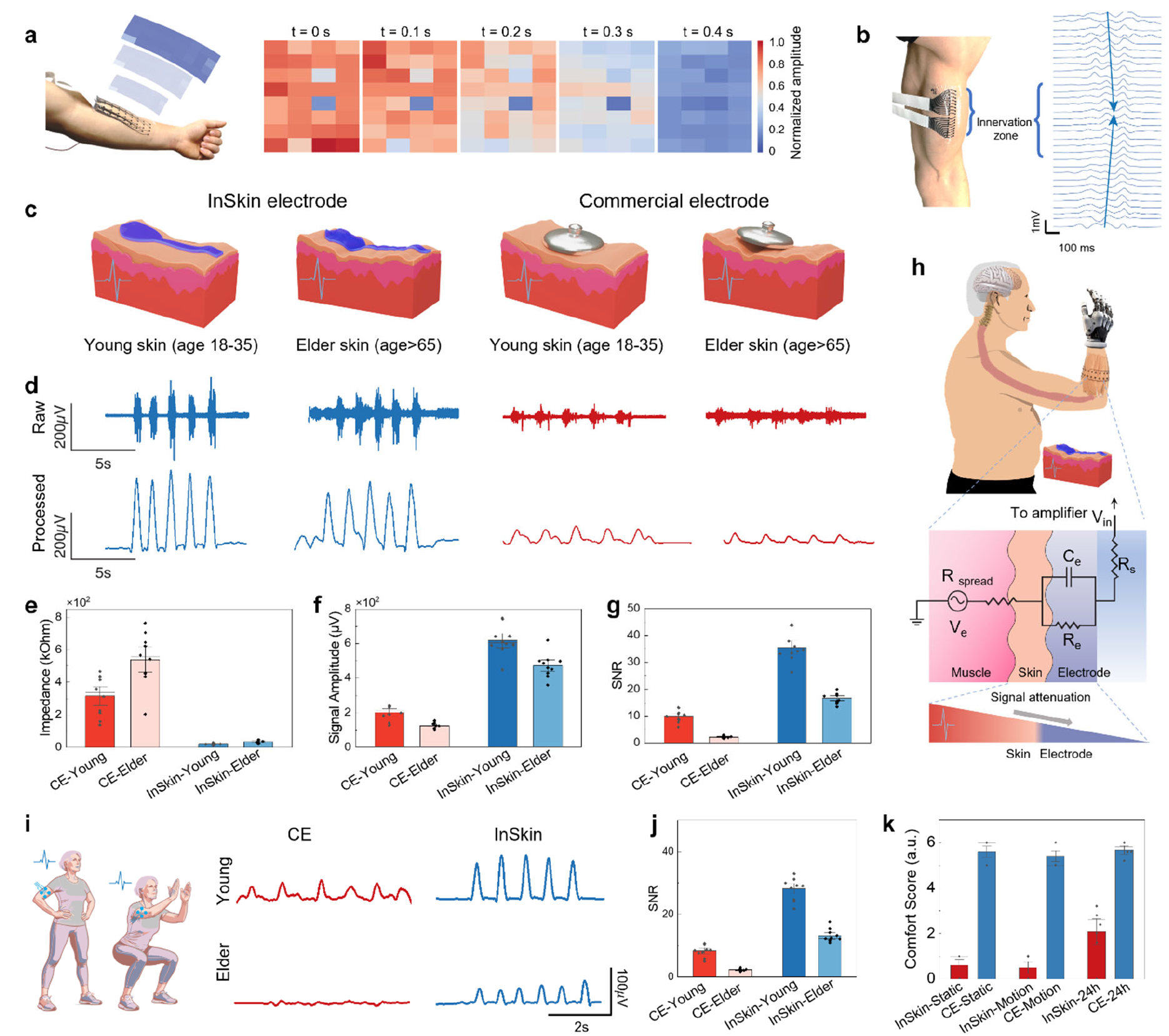
sEMG signal recording and group study of volunteers with different ages and skin conditions. (**A**) sEMG mapping of the forearm with a 32-channel array showing the change in normalized signal amplitude for each channel in heatmaps constructed from signals recorded each 100ms. (**B**) sEMG mapping using the 32-channel AdapSkin array for innervation zone localization on the bicep brachii. (**C**) Illustration of the electrode configurations employed to record sEMG signals, showing their different capability to conform to the skin’s surface features. (**D**) Representative sEMG signals of the repeated flexion of the bicep brachialis muscle recorded with CE and AdapSkin electrodes from a young volunteer and an older volunteer. (**E-G**), Bar graphs showing the impedance (F), absolute signal amplitude (F), and signal-to-noise ratio (G) for sEMG signals recording by CE and AdapSkin electrodes on Young and Elder groups (n.s. not significant). (**H**) Schematic diagram highlighting the importance of skin-electrode interface for signal attenuation. **(I)** Representative sEMG signals of the repeated flexion of the bicep brachialis muscle recorded with CE and AdapSkin electrodes from a young volunteer and an older volunteer when doing squats. (**J**) Bar graph showing the signal-to-noise ratio during the motion for the CE and AdapSkin sEMG recordings from the Young and Elder groups (n.s. not significant). (**K**) User comfort scores for different electrodes during static, dynamic, and long-term wearing conditions, with statistical significance marked.

To evaluate the performance of AdapSkin for sEMG recording across different age-related skin conditions, human subjects with two different age groups (the Elder group with ages ranging from 65 to 85 and the Young group with ages from 18 to 35) are recruited for sEMG measurements following an authorized IRB protocol (Fig. 3C). sEMG signals from the bicep brachialis muscle under repeated flexion are recorded in a bipolar configuration using both AdapSkin electrodes and CEs of the same size. The representative sEMG recordings are shown in Fig. 3D, indicating a higher signal quality is obtained from the AdapSkin recordings. Data analysis across the two study groups (10 subjects in each group) shows that the measured skin impedance increased ∼70.1% from the Young group to the Elder group when recording using the CE, while an increase of only 14% was observed when using AdapSkin (Fig. 3E). Consequently, the signal amplitude and SNR reduced 47%, and 80% respectively, from the Young group to the Elder group with the same muscle action when recording using the CE, compared to 22.5% and 42.6% when using the AdapSkin (Figs. 3F-H). Overall, our results show that the AdapSkin electrodes do not only decrease the skin-electrode impedance but, more critically, can lead to much-reduced changes in electrode-skin impedance, signal amplitude, and SNR across age groups (Fig. 3I), demonstrating its ability to provide reliable signals and fair recording across varying skin conditions. Conventionally, the reduced EMG amplitude is regarded as age-related sarcopenia (muscle loss) or motor neuron degradation (*34,35)*. In contrast, our results indicate that skin aging, and importantly the skin-electronic interface, all play very significant roles in signal attenuation for elderly people, while designing a better interface would help reduce such age-induced discrepancy. To evaluate the performance of the AdapSkin during motion, bicep contractions in dynamic (doing squat) conditions were evaluated, and it was shown that the SNR reduced by 75.3% when using CE electrodes, while only 17.1% SNR reduction was observed when using AdapSkin electrodes. Our user study also showed that AdapSkin offers improved comfort for short and long wear periods (Fig. 3J). Assessing the electrodes’ performance between hairy and hairless (shaved) skin among the participants of the Young group also revealed that the AdapSkin electrodes made by CP-G 250 showed reduced changes in electrode-skin impedance, signal amplitude, and SNR, when compared with the commercial electrodes (fig. S10).

### Robotic Control and Gesture Recognition

Gesture recognition through sEMG recordings is one of the most important techniques in developing human-machine interface (HMI) systems. These capabilities enable neural control of prosthetic devices for individuals with disabilities and facilitate robot control in both real-life and virtual-reality (VR) environments (*9*–*11*). We evaluated the performance of the AdapSkin array for gesture recognition by using the multi-channel sEMG recording to control a robotic hand with five independent fingers. Fig. 4A shows a gesture recognition system diagram, consisting of a multi-channel signal acquisition system, a computer for implementing the machine learning algorithm, the robotic hand, and the associated microcontroller module. The machine learning algorithm establishes a relationship between the subject’s gestures and the input signals, allowing the recognition and classification of hand movements using the offline sEMG dataset. The robotic hand then mimics the motion of the subject’s hand in real-time, enabling seamless gesture recognition and control. To demonstrate real-time gesture-to-prosthetic translation, the dataset including 10 distinct hand gestures was chosen to simulate the multiple conditions of the subject’s hand, by collecting the voltage signals captured from 32-channel sEMG sensors over the desired position of the subjects’ arm, aligning the first two sensing electrodes over the brachioradialis and then surrounding the forearm circumferentially (fig. S9). Fig. 4B illustrates the sEMG analog signals acquired at different stages of hand gestures, where all five fingers are moved independently. The electrical signals from the 32 independent sensors are captured and combined into a matrix that correlates with specific hand gestures. sEMG signals were acquired from an electrode array aligned over the elbow crease to maintain consistent landmark-based sensor placement (fig. S11). A convolutional neural network (CNN)-based machine learning framework is used for learning these signals (see details in Methods and Materials section). The corresponding voltage profiles for each gesture highlight the unique signal patterns associated with each movement (Fig. 4C). When the training process of offline sEMG dataset is finished and real-time gesture-to-prosthetic starts, the sEMG recordings from the same participant will be wirelessly transmitted to the computer and processed using the same setting (including a conventional sEMG processing bandpass filter (50-500 Hz), normalization and passed through a custom feature extraction algorithm (fig. S11). These processed signals serve as input features for the trained model, which predicts the corresponding hand gesture and sends control signals to the robotic hand controller continuously. The real-time gesture recognition algorithm operates using a 1-second sliding window, converging to a defined gesture when a match is detected with the training data at a confidence level exceeding 95% (Supplementary Movie 2). The confusion matrix (Fig. 4D) illustrates the classification results, where 7 of the 10 gestures achieved an accuracy greater than 98%, and the overall accuracy was 97.7%. The results provided evidence of the platform’s potential for non-invasive body-machine interfaces, which could significantly improve the quality of life for individuals with limb loss or mobility impairments.

**Fig. 4.**
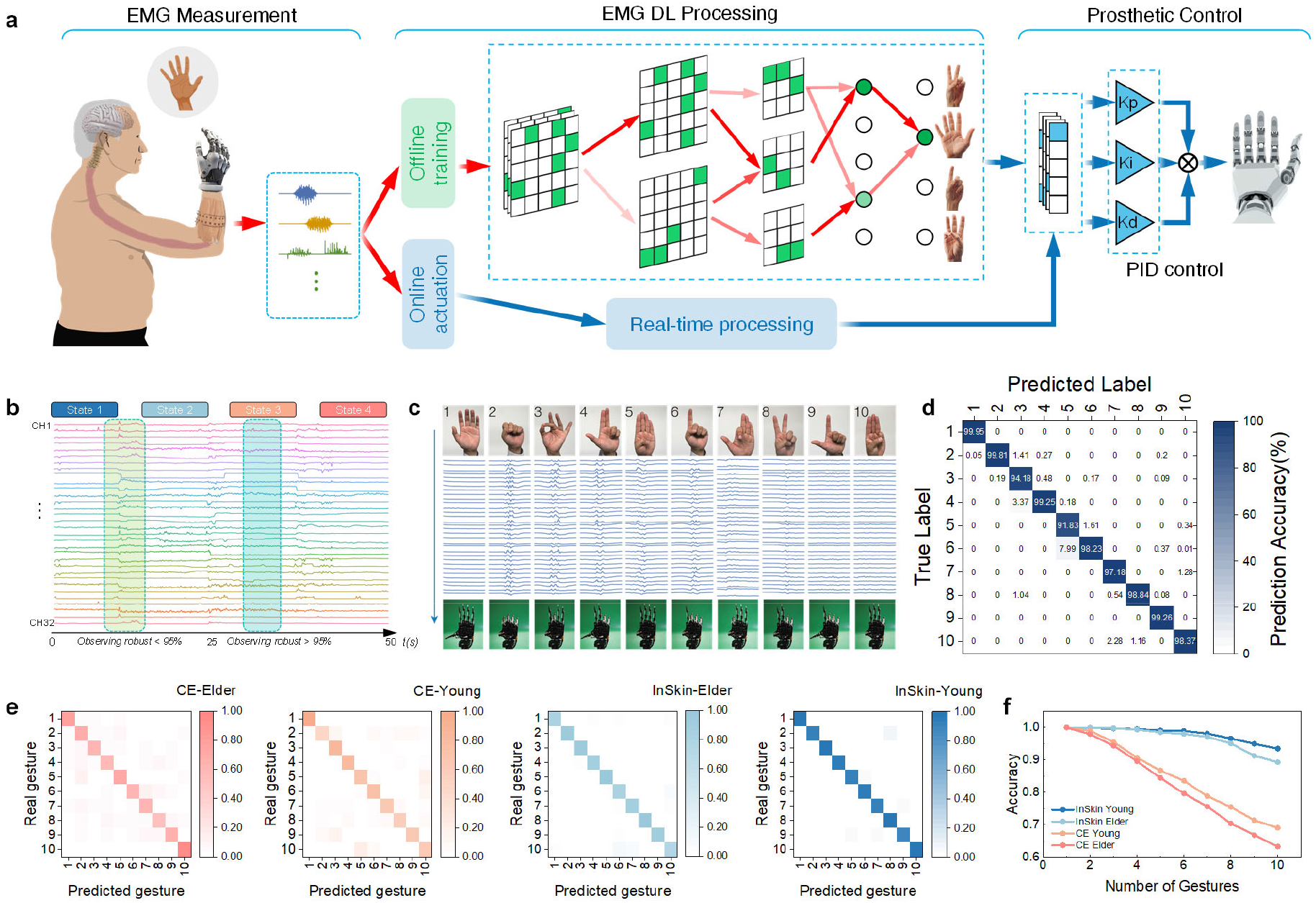
AdapSkin sEMG recording for gesture recognition and robotic hand control in human-machine interface systems. (**A**) Deep learning (DL)-assisted robotic control process flow, including sEMG data acquisition, DL model training process, and real-time robotic hand actuation. (**B**) Example of sEMG analog signals recorded from all 32 channels during finger motion. The continuum trial contains different gesture characteristics, including the hold period and transition period. The signal analysis shows that during the hold period, the recognition result of the gesture has the highest accuracy, while in the transition period between two different gestures, the recognition result is decreased beyond a threshold value. sEMG signals were acquired from an electrode array aligned over the elbow crease to maintain consistent landmark-based sensor placement (fig. S11). (**C**) Representative sEMG voltage profiles for 10 gestures and their signal patterns associated with each movement in real-time sEMG recording and control experiment. (**D**) Classification confusion matrix of 10 gestures recognition experiment through 32-channel AdapSkin recording. Blue and white text values are percentages of correct and incorrect predictions, respectively. (**E**) Comparison of performance differences between AdapSkin and commercial electrodes in the Young and Elder groups through 6-channel sEMG recording. (**F**) Data analysis shows the relationship between the number of gestures and accuracy when using AdapSkin and commercial electrodes to test the Young and Elder groups.

To compare the performance of AdapSkin and commercial electrodes across varying age-related skin conditions, the Elder and Young participants were asked to perform the multiple-gesture training and recognition experiment following the established protocol (Fig. 4E). To make a fair comparison, 6-channel recordings are used for both AdapSkin and CE electrodes due to the large size of the CE electrodes (fig. S11), while same training process and algorithm are used for the prediction. The representative results demonstrate a significantly higher signal quality from the AdapSkin recordings in both groups. Data analysis reveals that AdapSkin achieves recognition accuracies of over 95% for the elderly and over 97% for younger individuals, whereas CE recordings result only exceeding 60% for the elderly and 70% for younger participants in the 10-gesture recognition experiment. The relationship between the number of gestures and accuracy for both AdapSkin and CE recordings in both the Young and Elder groups is shown in Fig. 4F. Overall, our results demonstrate that data quality plays a vital role in the prediction accuracy when the same training and computational processes are applied, while the AdapSkin could lead to a very fair interface for high-fidelity data recording for both the Young and Elder groups, indicating its potential to create more fair and inclusive neural interfaces.

## Discussion

The AdapSkin platform represents a critical advancement in bioelectronics for surface electromyography and adaptive human-machine interfaces. By developing highly soft, stretchable, and conformable electrodes from conductive polymer composites, we have created a wearable sEMG array that bonds seamlessly to varied skin topographies and conditions across different age groups. The key innovations of the AdapSkin platform include dramatically reducing data variation and inequality of skin-electrode impedance and significantly improving the signal-to-noise ratio and sEMG amplitude compared to commercial rigid electrodes. The high spatial resolution of the sEMG arrays enables long-term mapping of complex neuromuscular activity and muscle innervation zones, offering unprecedented insights into muscle function and aging.

Our experiments particularly highlight the critical role of interface design for future human-machine interfaces, as such designs ensure high-quality data acquisition across diverse populations. This, in turn, significantly improves the accuracy of future machine intelligence, which relies heavily on advanced algorithms, abundant data, long-term training, and computing power. As global populations age, the demand for equitable, personalized bioelectronic solutions will continue to rise. Looking ahead, the intrinsic softness, stretchability, and customizability of the AdapSkin platform open numerous exciting applications in prosthetics, rehabilitation, neuromuscular disease monitoring, and injury prevention. We also expect such adaptive bioelectronic solutions to advance the future of AR/VR designs, which are often not designed with senior individuals in mind.

## Supporting information

Video S1. HD-sEMG mapping

Video S2. ML-enabled gesture recognition and wireless prosthetic control

## Data availability

The datasets generated during and/or analyzed in this study are available from the corresponding author upon reasonable request. The source data are provided in this paper.

## Ethical Considerations

### Study Design and Objectives

The validation of a novel high-density and stretchable surface EMG array guided the inception of this research. The primary aim was to explore the potential of flexible and stretchable electrodes for neuromuscular signal detection, heralding new horizons in rehabilitation tools and strategies.

### Recruitment and Informed Consent

The recruitment process was meticulously planned to encompass a diverse subject pool, including Michigan State University students, faculty, staff, and elderly individuals recruited through ResearchMatch.org. Participants were provided with a comprehensive consent form, ensuring complete transparency and voluntary participation, reflecting the inclusivity and accessibility objectives of the research.

### Procedural Integrity and Safeguards

The study’s procedures were designed with the utmost consideration for participant comfort and safety, employing sEMG sensors made from biocompatible materials for a non-invasive approach. Privacy and confidentiality were paramount, with rigorous measures such as randomly generated ID numbers and secure data storage.

### Potential Risks, Benefits, and Compensation

A thorough risk assessment concluded that there were no foreseeable risks to participants. While there were no direct benefits, the findings may significantly benefit the broader scientific community. Participants were compensated with a $20 Meijer gift card, acknowledging their valuable contribution.

### Compliance, Privacy, and Confidentiality

Adherence to ethical guidelines was unwavering, with compliance with equitable subject selection, adequate confidentiality provisions, and safeguards against coercion or undue influence. The privacy of subjects was stringently protected through an organized system, ensuring segregation of responsibilities and secure data handling.

## Acknowledgments

Author contributions: J.L. conceived the idea and directed the project. V.M. and J.L. designed the experiments. V.M. synthesized conductive composites. V.M., L.X., and C.M. designed and fabricated the device and performed the mechanical, electrical, and electrochemical measurements. V.M. performed the SEM. V.M., L.X., C.M., X.Y. conducted the sEMG experiments and human studies. J.W. contributed to the initial materials selection. V.M. developed the human study and protocol. C.X.C. and S.I. helped to prepare the schematics. Z.T. helped to analyze the sEMG signals. L.X., V.M., J.D., and C.M. wrote the custom algorithm for signal processing and ML-enabled gesture recognition. V.M., L.X., and J.L. wrote the manuscript. All authors discussed the results and commented on the manuscript. All authors have approved the final version of the manuscript. The authors would like to thank Nancy Campbell and 19 other volunteers for their participation in the study.

