## Supplementary Information for "Highly Adaptive Conductive Polymer Electronics Enhance Neural Data and Learning Accuracy"

### Methods

### Materials

An aqueous dispersion of poly(3,4-ethylenedioxythiophene) polystyrene sulfonate (PEDOT:PSS) (CleviosTM PH1000; solid content ≈ 1.1−1.3 wt%; weight ratio of PEDOT to PSS ≈1:2.5; pH~2) was purchased from Heraeus. A WPU dispersion (ALBERDINGK® U4101; solid content ≈ 39−41 wt%; elongation at break~1400%; pH ≈ 7.0−8.5) was obtained from Alberdingk Boley. ALBERDINGK® U4101 is an aqueous, anionic dispersion of an aliphatic polyether-polyurethane without free isocyanate groups. 1-Ethyl-3-methylimidazolium ethyl sulfate (EMIM:ESO4) was purchased from Sigma-Aldrich. Glycerol and an ammonia hydroxide solution of 25% were purchased from Sigma-Aldrich. Polyurethane and adhesive acrylic double-layer (Artificial skin, AS) was purchased from 3M™. All chemicals and materials were used as received without further purification.

### Materials preparation and film deposition

To prepare the composite material for the sEMG array biosensor, the ionic liquid (EMIM:$ESO_4$) was diluted to 1.48 wt% with deionized water before being added to PEDOT:PSS dispersions. The ionic liquid solution was added dropwise to the PEDOT:PSS dispersion in a ratio of 15:25, and the mixture was stirred at 400 rpm for 12 hours. Before adding a waterborne polyurethane (WPU) dispersion, 0.15 wt% of an ammonia solution was added with respect to the PEDOT:PSS dispersion. The WPU was then added to the solution to obtain a final ratio of 15:25:85 PEDOT:PSS:IL:WPU, and the entire mixture was then stirred for 2 hours. This solution was named "Solution CP."

Solution CP-G was prepared by adding varying concentrations of glycerol to Solution CP, with ratios ranging from 1:1 to 1:300. Among the tested ratios, Solution CP-G 1:250 exhibited the most favorable properties for the intended task and was therefore selected.

The dispersions were then spin-coated or drop-casted onto bare or Polytetrafluoroethylene (PTFE)-coated glass slides and dried in a vacuum desiccator at room temperature overnight.

### Device Fabrication

The fabrication process of the InSkin is based on the vertical integration of stretchable functional layers and micro-patterning with direct laser engraving, as illustrated in **Fig. S1 A**. Glass slides (Fisherbrand™ Superfrost™ Plus Microscope Slides) coated with a chemically resistant polytetrafluoroethylene film (McMaster-Carr Chemical-Resistant Slippery Film Made from Teflon® PTFE) was employed as the initial substrate.

The as-prepared Solution CP-G was then drop-casted on the hydrophobic surface, and the solvent was evaporated overnight to create a uniform and smooth thin film. The surface of the Solution CP-G thin film was cleaned and lastly treated with oxygen plasma (Plasma Etch Inc. Model PE-50, 120 VAC 60Hz, 200 mTorr) for 5 minutes to improve the adhesion with the AS.

A Universal Laser System™ VLS6.75 with a $CO_2$ laser and High-Power Density Focusing Optics (HPDFO, a lens with a focal point of 0.001 inch (25.4 μm) at a power of 1.5 W and a speed of 11% was employed to create the pattern on the dried and plasma-treated Solution CP-G thin film. After laser patterning, the excess film is removed from the substrate, and the electrode and interconnect pattern are released. Upon pattern release, an AS rectangle with the coating protecting the adhesive layer removed is aligned and attached to the patterned Solution CP-G to transfer the electrodes onto the substrate. The transferred array is then treated a second time under oxygen plasma to improve the adhesion with the encapsulation layer and to improve the electrode interface to the skin. After plasma etching, the array is connected with flat-flexible cables through an anisotropic conductive tape before annealing at 80℃ for 15 minutes to improve the electrical connection. A second AS layer is laser patterned to create openings corresponding to the electrode pads on the array to encapsulate the device. The patterned AS is then aligned and applied on the connected array to isolate the interconnects while leaving exposed sensing pads.

After encapsulation, a solution of PEDOT:PSS:Glycerol in a 1:1 ratio named Solution P is drop-casted on each sensing electrode to create a softer interface with the skin. The device is then annealed one last time at 60°C for 10 minutes before being applied to the skin.

### Human Studies:

To evaluate the array's performance on diverse skin morphologies, we recruited 20 participants through campus advertisements and *ResearchMatch.org*. The cohort ranged in age from 19 to 83 years, with a gender balance of 50% male/50% female. Interested respondents received detailed information and were scheduled for sEMG recording sessions. The Institutional Review Board at Michigan State University approved the study procedures under reference number 00008023. All participants completed written informed consent before participation. Anonymization utilized randomly generated identification numbers, and all data were securely stored in encrypted formats.

We adhered closely to Michigan State University's guidelines for participant selection. Using the selected participants, we used the three electrode types (CE, Solution CP, and Solution CP-G) to monitor the sEMG signals of the biceps brachii. The biceps brachii was chosen as the muscle to be tested due to its relatively large area and the simplicity of its flexion.

The signal-to-noise ratio (SNR) was quantified as the ratio between the steady-state sEMG signal amplitude during sustained contraction and the standard deviation of the baseline noise during rest. SNR values were compared between commercial, Solution CP, and Solution CP-G electrodes using paired t-tests.

### Comfort Study

To comprehensively assess the electrodes' comfort under **static**, **dynamic**, and **chronic** conditions, we assessed participant comfort while wearing the electrodes using a structured questionnaire. The questionnaire was designed to evaluate comfort during periods of stillness (static), movement (dynamic), and prolonged wear (chronic). Participants rated their experience with two electrode

types: a dry electrode and the Solution CP-G 250 electrode, providing feedback on comfort, discomfort, and overall preference for extended use.
Twenty participants, aged 19 to 83, were recruited for the study. After each recording session, they were asked to complete the questionnaire, which covered the following categories:

- **Static condition**: Comfort while seated or stationary.
- **Dynamic condition**: Comfort while moving or performing mild physical activity.
- **Chronic condition**: Comfort and preference for long-term electrode wear, including during sleep or extended daily use.

The following table summarizes the questions:

| **Category** | **Question** | **Response Options** |
|---|---|---|
| **Static Comfort** | How comfortable are you wearing the first electrode (dry) while seated or stationary? | Very comfortable, Somewhat comfortable, Neutral, Somewhat uncomfortable, Not comfortable |
| | How comfortable are you wearing the second electrode (CP-G) while seated or stationary? | Very comfortable, Somewhat comfortable, Neutral, Somewhat uncomfortable, Not comfortable |
| **Static Discomfort** | How often do you experience discomfort while wearing the first electrode (dry) when stationary? | Never, Rarely, Sometimes, Often, Always |
| | How often do you experience discomfort while wearing the second electrode (CP-G) when stationary? | Never, Rarely, Sometimes, Often, Always |
| **Dynamic Comfort** | How comfortable are you moving around while wearing the first electrode (dry)? | Very comfortable, Somewhat comfortable, Neutral, Somewhat uncomfortable, Not comfortable |
| | How comfortable are you moving around while wearing the second electrode (CP-G)? | Very comfortable, Somewhat comfortable, Neutral, Somewhat uncomfortable, Not comfortable |
| **Dynamic Discomfort** | How often do you experience discomfort while moving with the first electrode (dry)? | Never, Rarely, Sometimes, Often, Always |
| | How often do you experience discomfort while moving with the second electrode (CP-G)? | Never, Rarely, Sometimes, Often, Always |
| **Chronic Wear** | How comfortable would you feel wearing the first electrode (dry) for longer than one day? | Very comfortable, Somewhat comfortable, Neutral, Somewhat uncomfortable, Not comfortable |

| | | |
|---|---|---|
| | How comfortable would you feel wearing the second electrode (CP-G) for longer than one day? | Very comfortable, Somewhat comfortable, Neutral, Somewhat uncomfortable, Not comfortable |
| **Preference** | Which electrode would you prefer to wear for long periods (more than one day)? | Dry electrode, CP-G electrode |
| **Sleep Impact** | How comfortable would you feel wearing the first electrode (dry) while sleeping? | Very comfortable, Somewhat comfortable, Neutral, Somewhat uncomfortable, Not comfortable |
| | How comfortable would you feel wearing the second electrode (CP-G) while sleeping? | Very comfortable, Somewhat comfortable, Neutral, Somewhat uncomfortable, Not comfortable |
| **Skin Irritation** | How often did you experience any skin irritation with the first electrode (dry)? | Never, Rarely, Sometimes, Often, Always |
| | How often did you experience any skin irritation with the second electrode (CP-G)? | Never, Rarely, Sometimes, Often, Always |

**Data Analysis**

Comfort scores were analyzed using repeated measures ANOVA, with post-hoc Tukey correction, to determine significant differences between the two electrode types under static, dynamic, and chronic conditions. These ratings provided insights into participant preferences, skin irritation, and overall comfort, especially during periods of movement and extended wear.

**Biopotentials recording**

**Surface electromyography (sEMG):**

The sEMG signals were recorded by an INTAN RHD recording controller (intan TECHNOLOGIES™) interfaced to a 32-channel amplifier chip RHD recording headstage (intan TECHNOLOGIES™). The proprietary software controlling the amplifier (RHX Data Acquisition Software, intan TECHNOLOGIES™) can produce spectrograms and Inter-spike-interval histograms during data collection to confirm signal detection, as shown in **Fig. S7 B, C**. After application of the sensing electrodes on the subject's skin, the electrodes were connected to the recording headstage through a custom PCB connector that could be connected to the sEMG array through the FFC connectors. The sEMG signal was recorded with a sampling rate of 2.5 kHz, a high-pass filter of 10 Hz, a low-pass filter of 500 Hz, and a notch filter of 60 Hz and its harmonics. To obtain sEMG recordings, the reference electrode was placed on the skin, far from the recorded muscle and preferably on a bony prominence, while the two recording electrodes were placed 1 cm apart on the muscle being tested when recording single channel sEMG signals. Measurement recording was started, and bicep testing was performed as a sample motion. For the bicep testing, the reference electrode was placed on the top of the hand, and the two working electrodes were placed 1 cm apart on the bicep brachii symmetrically with respect to the peak of the muscle upon flexion. A representative sEMG signal composed of 10 different muscular contractions recorded with a 32-channel array is shown in **Fig. S7 D**.

### Electrocardiogram (EKG):

The EKG signals were recorded using a 3-electrode configuration and using the same setup during sEMG data collection. After applying the sensing electrodes on the subject's skin, the electrodes were connected to the recording head stage through a custom PCB connector that could be connected to the EKG electrodes through a wired adapter. The EKG signal was recorded with a sampling rate of 2.5 kHz, a high-pass filter of 10 Hz, and a low-pass filter of 200 Hz. A representative EKG trace with clear P and T waves and QRS complex is shown in **Fig. S7 E**.

### Electroencephalogram (EEG):

The EEG signals for alpha wave detection were recorded using a 3-electrode configuration by placing two sensing electrodes in locations FP1 and FP2 and by placing the reference electrode in location O2 on the scalp of the tested subjects. The same setup employed during the sEMG data collection was used to record EEG signals. After applying the sensing electrodes on the subject's skin, the electrodes were connected to the recording head stage through a custom PCB connector that could be connected to the EKG electrodes through a wired adapter. The EEG signal was recorded with a sampling rate of 2.5 kHz, a high-pass filter of 1 Hz, and a low-pass filter of 100 Hz to minimize noise. After baseline recording, the subjects were asked to close their eyes and breathe deeply so as to induce a relaxed state and detect alpha brain waves. A representative EEG trace before and after breathing exercises is shown in **Fig. S7 F**, where the 10 Hz peak, representative of alpha waves, is clearly seen.

### Robotic Hand Control

#### Classification

Ten hand gestures were selected for analysis (as "configuration" is intended, the finger number engaged in the movement with respect to the relaxed configuration): 1: thumb, 2: index, 3: middle finger, 4: ring finger, 5: pinky.: hand relaxed, hand closed in a fist (12345 fingers configuration), thumb and middle finger adduction (13 fingers configuration), thumb, index, and middle finger relaxed (45 fingers configuration), thumb opposed (1 fingers configuration), index alone (1345 fingers configuration), index contracted (2 fingers configuration), peace sign (145 fingers configuration), "L" sign (345 fingers configuration), thumb-pinky touching (15 fingers configuration). Surface electromyography (sEMG) signals were acquired from an electrode array aligned over the elbow crease to maintain consistent landmark-based sensor placement (**Fig. S7 A**).

Participants attended two data collection sessions on separate days to furnish training and test data sets for the gesture classification model. The classification pipeline comprised preprocessing, feature extraction, and classification stages. Raw sEMG data were recorded using an INTAN acquisition system, and bandpass was filtered to remove noise, offsets, and motion artifacts. The recording instrument can display in real time the signal spectrogram (**Fig. S7 B**) and the inter-spike interval histogram (**Fig. S7 C**) to quickly confirm successful spike detection. Preprocessing included buffer deletion (0.1-0.25 s), normalization, and smoothing. Processed data were input to a pattern recognition neural network algorithm in Python 3.6, starting with three 3×3 filters as a convolution layer, followed by max pooling, three fully connected neural networks, and a dropout layer. ReLU activation was used for the convolution layer and two fully connected layers, while hyperbolic tangent sigmoid activation was used for the last fully connected layer. The final output comprised 10 neurons with SoftMax activation, representing the 10 gestures in this experiment. The dataset was split 80:20 for model training and testing.

To optimize sEMG signal amplitude, participants were instructed to exert 70-80% of their maximal voluntary contraction during data collection. To preclude fatigue, the 10 gestures could be acquired within 10 minutes. Participants could also pause and rest as needed.

**<u>Recognition</u>**

The recognition and robot hand control experiment followed the second session after adequate rest. sEMG data were sampled every 1 s based on the preceding 0.5 s recording window after preprocessing. A motion onset detection algorithm first screened for movement initiation by thresholding the mean absolute sEMG amplitude. If the threshold was not exceeded, the recognition output and robot hand state were maintained. Otherwise, the trained model classified the detected gesture, and the predicted label was mapped to servo motor commands to actuate the robot hand (**Fig. S7 I**). Participants completed all 10 gestures while holding each posture until the robot hand achieved the target configuration, regardless of classification accuracy. The full protocol was completed within 5-10 minutes to avoid fatigue.

**<u>Hardware Framework</u>**

The robot hand digits were driven by five individual servo motors, enabling independent flexion-extension of each finger via clockwise-counterclockwise rotations. A microcontroller (ATMega328P, Microchip Technology) managed servo command receipt, analog-digital conversion, and motor control. Gesture predictions from the PC were mapped to pulse width modulation (PWM) signals and sent to a 16 CH PWM servo motor driver (PCA9685, NXP Semiconductors) to operate the five motors.

**<u>Characterization</u>**

**<u>SEM:</u>** The SEM images were taken by a JEOL 6610LV SEM (15 kV).

**Mechanical Property Characterization:** The mechanical properties of freestanding Solution CP and WG and their substrate-supported films, approximately 100 µm in thickness, were studied using the CellScale UniVert with a 20-N loading cell. The samples were clamped between tensile testing grippers. A caliper measured the width and length of the film, while an optical microscope measured the thickness. The composite films had rectangular symmetry with a sample size of $30 \times 10 \times 0.1$ $mm^3$ (length × width × thickness). A single stretching step was performed at a stretching rate of 1%/s.

**Electrical and Electrochemical Characterization**

**Resistance Measurement:** To measure the resistance of the Solution CP and CO-G composites during stretching, samples of dimensions $10 \times 25 \times 0.1$ $mm^3$ were clamped to a UniVert testing system. Contacts were made by soldering copper wires to the strip using silver epoxy as the soldering paste. Alligator clips connected the copper wires to a high-resolution potentiostat (PalmSens4, PalmSens). Chronoamperometry was recorded for the duration of the elongation test with a 0.5V bias applied to the conductive material.

**Electrochemical Impedance Spectroscopy (EIS):** EIS measurements were performed with the same high-resolution potentiostat, and the impedance was recorded from 10 MHz to 10 Hz in the PBS buffer solution (pH 7.4). The specific settings included a sine wave frequency from 1 Hz to 1 MHz and a signal amplitude of 10 mV.

**Cyclic Voltammetry (CV):** CV measurements were carried out by collecting and averaging 10 scans from -0.5V to +0.5V at a scan rate of 0.1V/s with the same high-resolution potentiostat. The scans were recorded in 1X phosphate-buffered-saline (PBS) buffer solution, and the enclosed area was computed to quantify the composite changes under elongation.

**Effects of Stretching:** Both electrode types showed a less than 4-fold increase in impedance under 100% elongation (**Fig. S6 C**). Figs. S6 C-G provide specific details about the effects of stretching on impedance and CV measurements.

**Thermal Stability:** Lastly, Solution CP-G 250 showed a 0.64% decrease in resistance when heated up to 45℃, comparable to the 0.16% decrease of a silver electrode, confirming the material stability in scenarios with conditions similar to their intended use (**Fig. S6 H**).

### Supplementary Figures

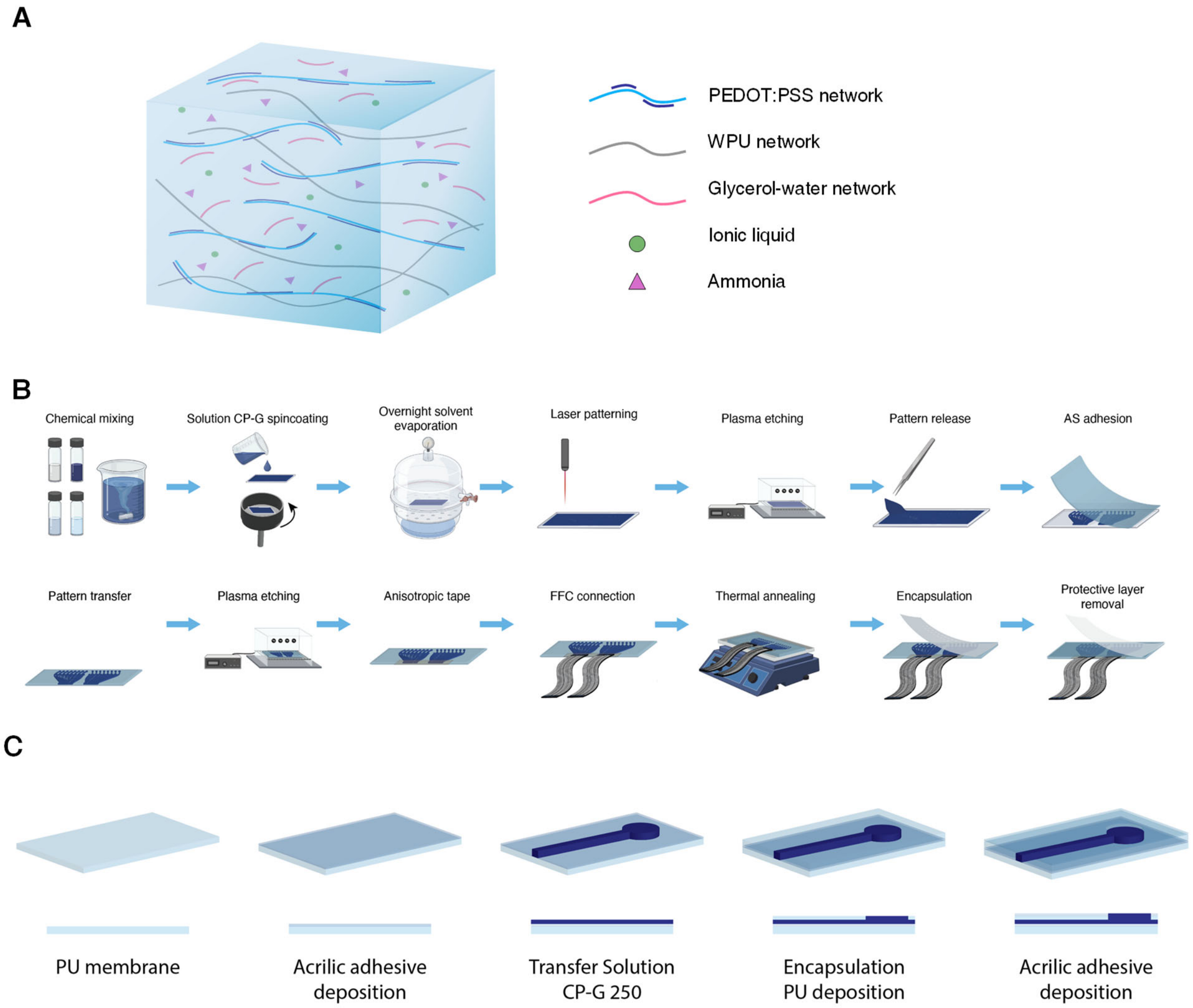


**Supplementary Figure S1 | A**. Schematic representation of the polymer composite structure with its chemical constituents. The network consists of PEDOT:PSS (blue), waterborne polyurethane (WPU) (gray), and a glycerol-water network (pink), with ionic liquid (EMIM:ESO$_4$) (green) and ammonia (purple) dispersed throughout. The final composition was prepared by mixing PEDOT:PSS, EMIM:ESO$_4$, WPU, and ammonia in a volume ratio of 124:186:23:1, respectively. PEDOT:PSS was diluted to 1.3% weight in $H_2O$, while EMIM:ESO$_4$ was diluted at 1.34% wt% in deionized water (DI water). WPU was used as purchased, with a solid content of 39–40%. **B**. Fabrication process for the multichannel high-density sEMG array for real-time prosthetic control: (1) poly(3,4-ethylenedioxythiophene) polystyrene sulfonate (PEDOT:PSS), 1-Ethyl-3-methylimidazolium ethyl sulfate (EMIM:ESO$_4$), water-borne polyurethane (WPU), and Glycerol were mixed during a 2-day process to obtain Solution CP-G 250, (2) Solution CP-G 250 was drop-casted (0.02 mL/cm$^2$) or spin-coated (2000 rpm, 30 seconds) on a polytetrafluoroethylene (PTFE)-coated glass slide to create a uniform thin film, (3) the coated slides were left to dry in a vacuum chamber overnight to allow for a faster solvent evaporation and humidity removal, (4) The dried films were patterned with a $CO_2$ laser (Power: 1.5 watts, speed: 11% of the instrument's maximum speed) to obtain flexible and stretchable sensing pads and interconnects, (5) prior to pattern release, the surface of Solution CP-G 250 was treated with oxygen plasma (5 min, 120 VAC 60Hz, 200 mTorr) to roughen the surface and promote adhesion with the artificial-skin (AS) substrate, (6) the excess Solution CP-G 250 not composing sensing pads or interconnects was released from the PTFE film to expose the patterned array, (7) a thin double layer of polyurethane and adhesive acrylic (artificial skin, AS) was overlayed on the patterned Solution CP-G 250 and adhesion was promoted by applying weights on the back side of AS, (8) AS was peeled off from the PTFE. The patterned Solution CP-G 250 was transferred to AS, and (9) the exposed surface was treated with oxygen plasma (5 min, 5 min, 120 VAC 60Hz, 200 mTorr) to improve the adhesion with the encapsulation layer and the impedance of the exposed sensing pads, (10) a strip of anisotropic tape was applied on the interconnects ends to allow for the electrical connection with the flat-flexible cables (FFC), (11) FFC cables were connected to the Solution CP-G 250 interconnects to interface the array with the recording instrumentation and prosthetic control system, (12) the FFC-connected array was thermally annealed at 80℃ for 15 minutes to improve the Solution CP-G 250-anisotropic tape-FFC cable interfaces and electrical connection, (13) a second layer of AS, patterned with openings corresponding to the electrodes pads, was applied on the Solution CP-G 250 array with the adhesive side facing as to allow for skin adhesion, (14) the AS's packaging and protective layer was removed to expose the AS's adhesive side. **C**, Perspective illustration of the multilayered device and the assembly sequence: (1) polyurethane membrane and (2) acrylic adhesive, which make up the bottom substrate, (3) Solution CP-G 250 to form sensing pads and interconnects, (4) polyurethane membrane and (5) acrylic adhesive to form the top encapsulation layer having the adhesive side facing the skin.

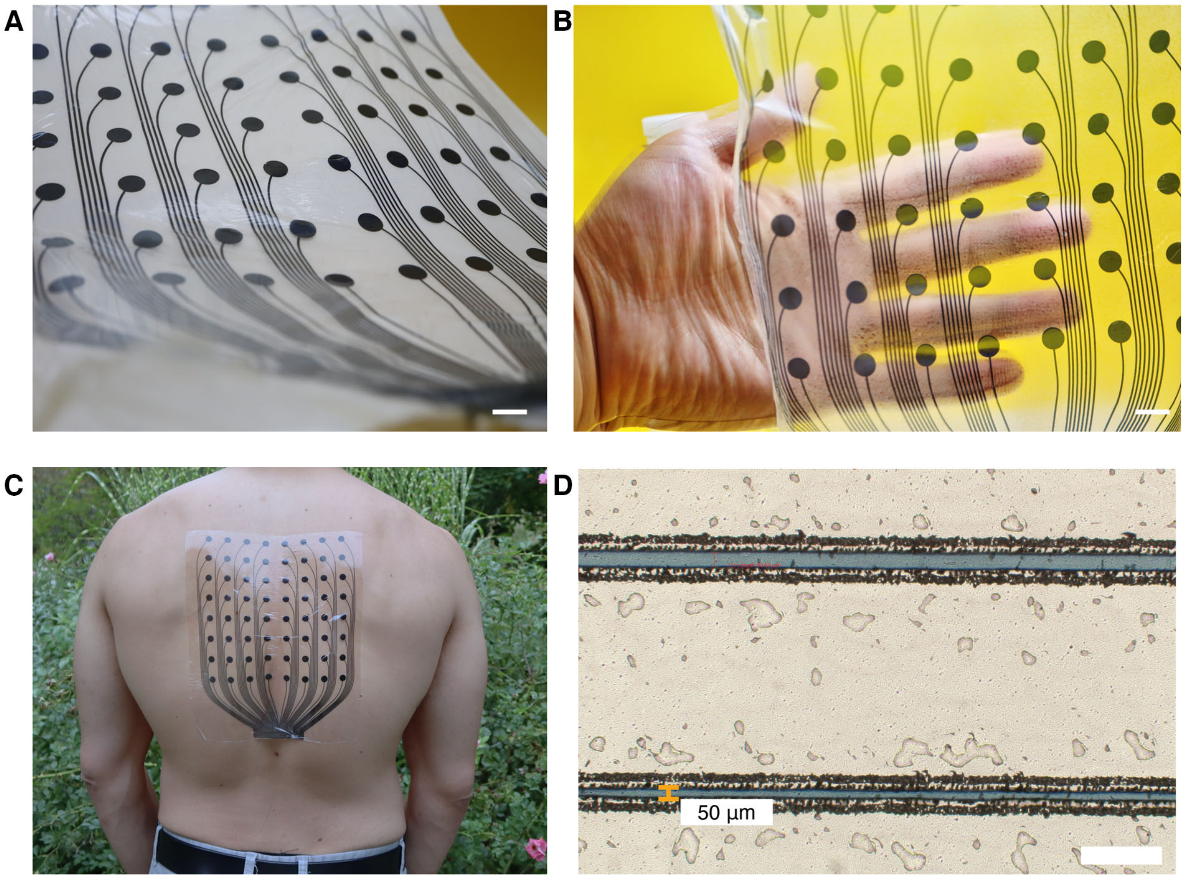


**Supplementary Figure S2 |** *Large-Area, High-Density sEMG Array*: **A**, **B,** Optical photographs of the flexible, transparent array illustrating its thin, stretchable design and close conformation to human skin. Scale bars: 2cm. **C**, Image showing the full-back 64-channel surface electromyography (sEMG) array conforming to the back, demonstrating its ability to cover large areas for muscle activity mapping. **D**, Microscope image showing the fine resolution of the sEMG array, with electrode interconnects spaced at 50 µm, highlighting the device's high precision and miniaturization capabilities. Scale bar: 200µm.

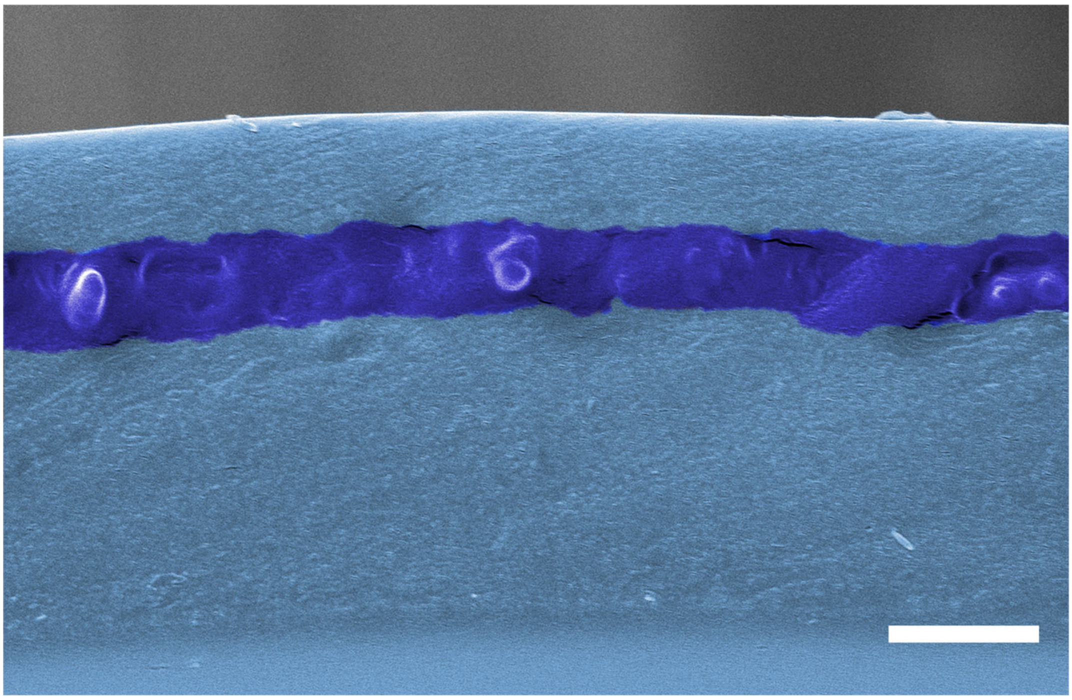

**Supplementary Figure S3 |** *Thickness of Solution CP-G 250 Arrays*: Pseudo-colored scanning electron microscope (SEM) image of the cross-section of Solution CP-G 250 encapsulated between two layers of AS, highlighting a multilayer structure with a total thickness of 33 μm. Scale bar: 10 μm.

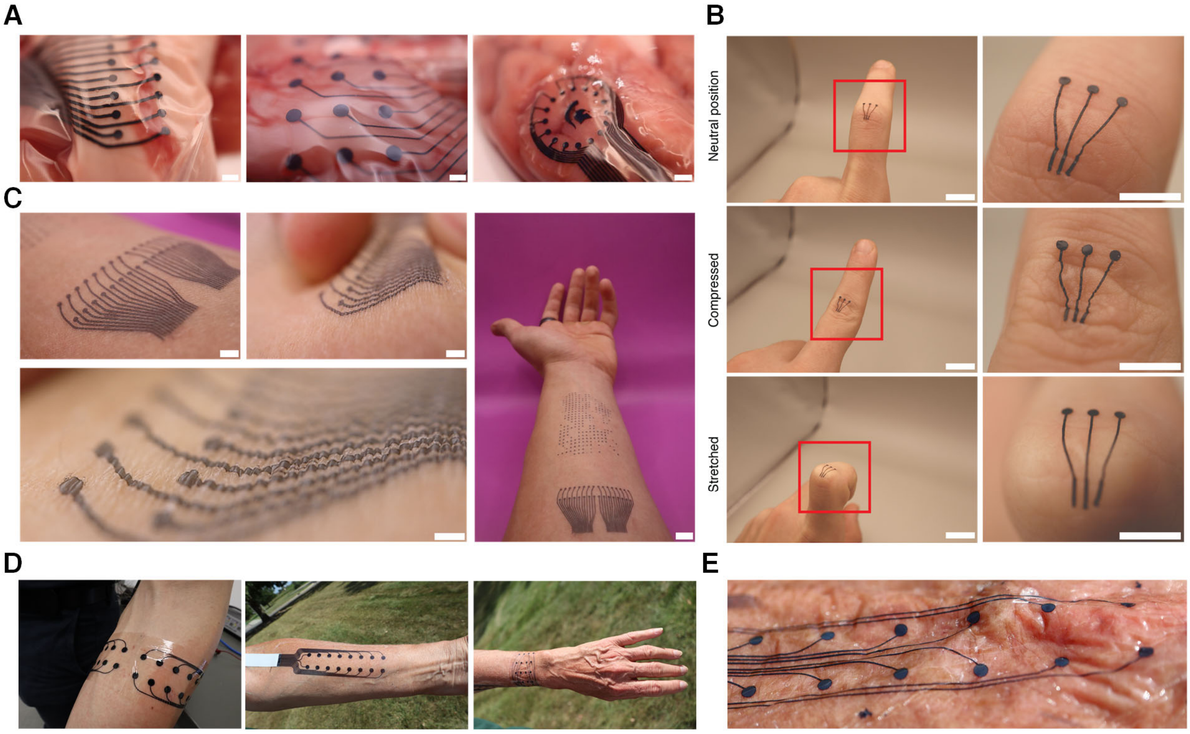


**Supplementary Figure S4** | *Flexibility, Conformability, and Patternability of InSkin Devices Made with Solution CP-G 250*: **A**, Photos of different devices applied on a pig brain ex-vivo. These images highlight the ability of the stretchable Solution CP-G 250 arrays to conform to soft tissue without detachment or loss of interface. Scale bars: 2 mm. **B**, Representative photos showing a 3-electrodes array conforming to the micro and macro features of the skin during stretching and compression. Left-side scale bars: 1 cm; right-side scale bars: 5 mm. A separate image showcases the patternability of Solution CP-G 250, with the iconic Spartan Helmet from Michigan State University patterned on the skin, demonstrating how sensors can be customized for patient comfort and aesthetics. Scale bar: 1 cm. **C**, Photographs of patterned Solution CP-G 250 applied on the skin, illustrating excellent conformability to skin features at different scales. Scale bars: 2 mm, 2 mm, 1 cm, 2 mm, and 1 mm, respectively. **D**, Photographs illustrating the conformability of the sEMG arrays on different body parts: forearm (left), forearm of an elderly subject (center), and wrist (right), demonstrating the flexibility and adaptability of the device to various skin textures and anatomical regions across both young and elderly subjects. **E**, A high-resolution 2x8 channel array adhering and conforming to the wrinkles of an elderly person's skin, showcasing the array's adaptability to uneven surfaces.

A

B

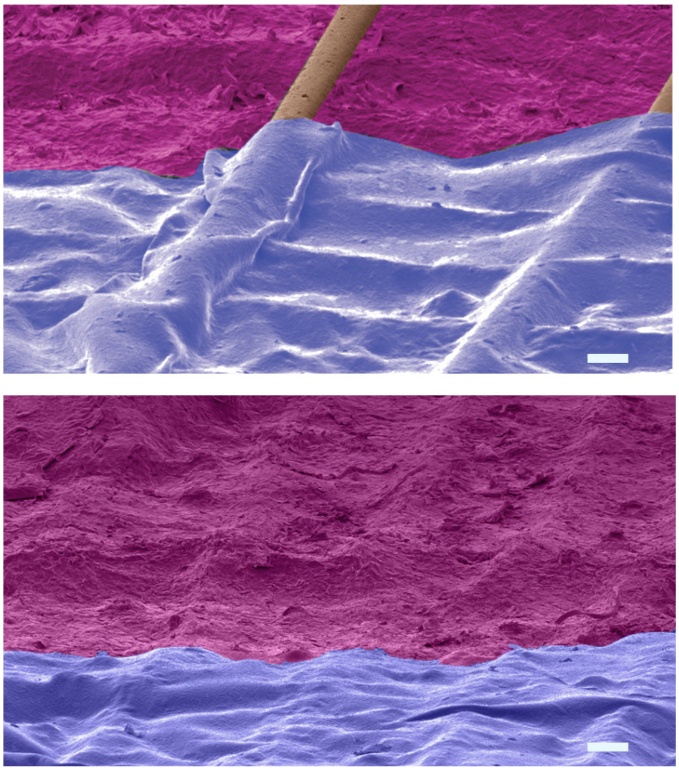

**Supplementary Figure S5 | A**, False color scanning electron microscope (SEM) image of a cross-section of Solution CP-G 250 on a wrinkled skin phantom with hairs, highlighting the material's thinness and surface conformity – scale bar 100 µm. **B,** False color scanning electron microscope (SEM) image of a cross-section of Solution CP-G 250 on a smooth skin phantom – scale bar 100 µm.

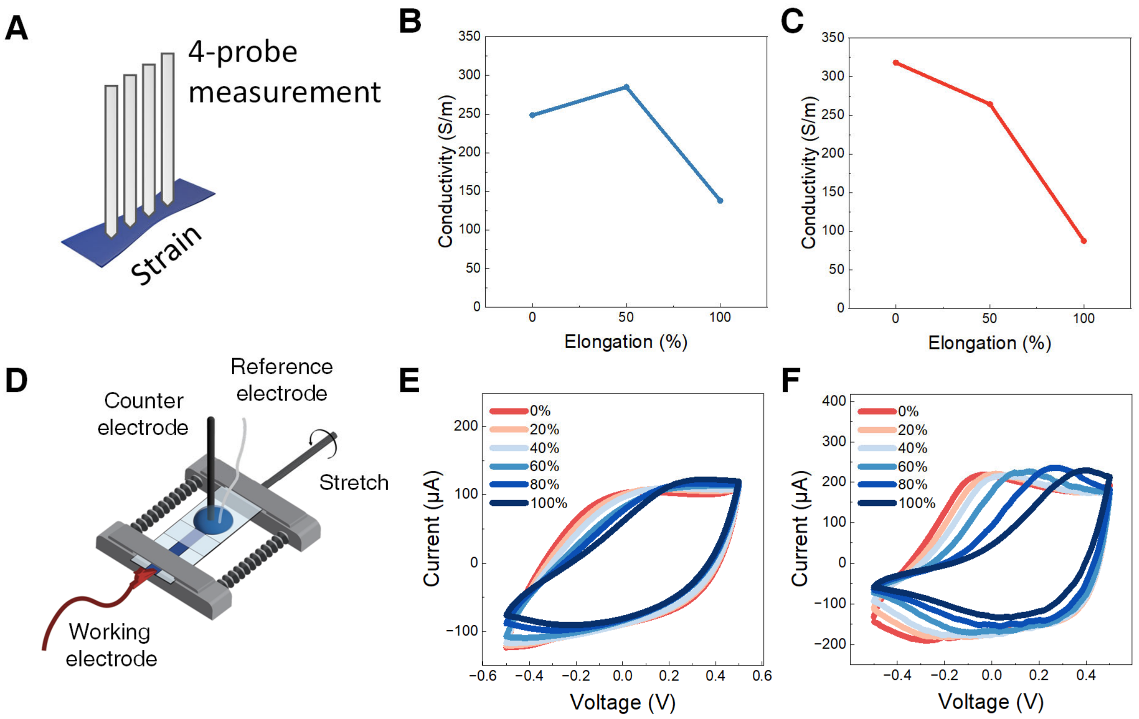


**Supplementary Figure S6 |** *Electrochemical and Mechanical Performance of Stretchable Composites:* **A,** Illustration of the experimental setups used for recording the electrical conductivity of the stretchable composite electrodes. **B, C,** conductivity as a function of elongation (0%, 50%, and 100%) for solution CP (**B**) and solution CP-G 250 (**C**). Both samples show a decrease in conductivity with increasing elongation, though solution CP-G 250 exhibits a more gradual reduction, indicating improved performance under mechanical strain. **D**, Illustration of the experimental setups used for recording electrochemical impedance, cyclic voltammetry, and force-strain characteristics of the stretchable composite electrodes. **E, F,** CV curves under varying stretching conditions (0% - 100%) for Solution CP (**E**) and Solution CP-G 250 (**F**), indicating consistent electrochemical behavior of both materials with increasing elongation, although Solution CP-G 250 maintains better performance under strain.

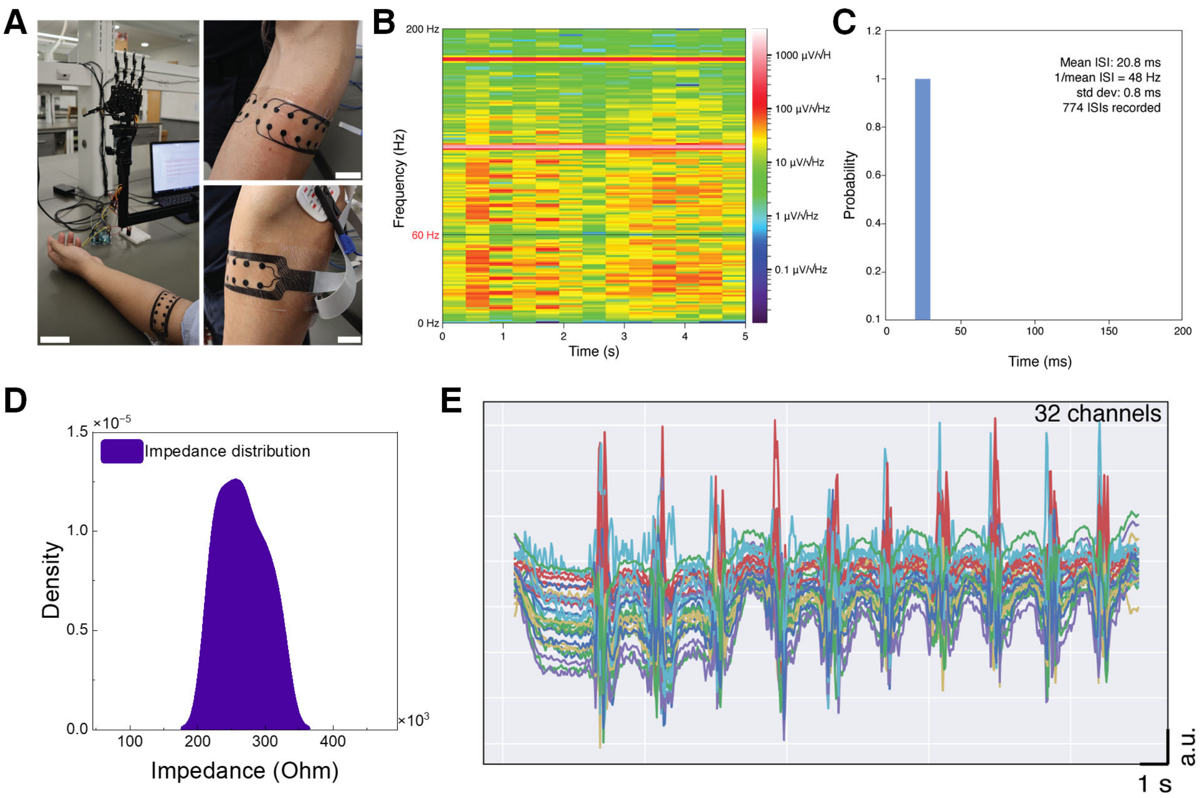


**Supplementary Figure S7 | A**, Photos of the prosthetic robotic hand control with the multi-channel HD-sEMG array. The array is wrapped around the subject's forearm and connected to a wireless amplifier (Mentalab LLC). Scale bars: left side: 5 cm, right side: 2 cm, **B**, Representative Spectrogram for an individual channel of the HD-sEMG array showing the power spectral density of the sEMG signal for different frequencies after a Fourier transform was applied to the signal. **C**, Representative Inter-Spike Interval (ISI) histogram for an individual channel of the HD-sEMG array, showing a probability = 1 of 774 spike events with a mean frequency of 48 Hz, mean inter-spike intervals of 20.8 ms, and a standard deviation for the ISI of 0.8 ms. **D,** Impedance density distribution for a 32-channel array interfaced with the skin, illustrating the narrow impedance range achieved using Solution CP-G 250, confirming the uniformity of the electrode-skin interface. **E**, Representative overlap of the sEMG signal recorded with a 32-channel sEMG array. The plot shows how the different channels behave similarly and perform thanks to the high composite uniformity and narrow impedance distribution.

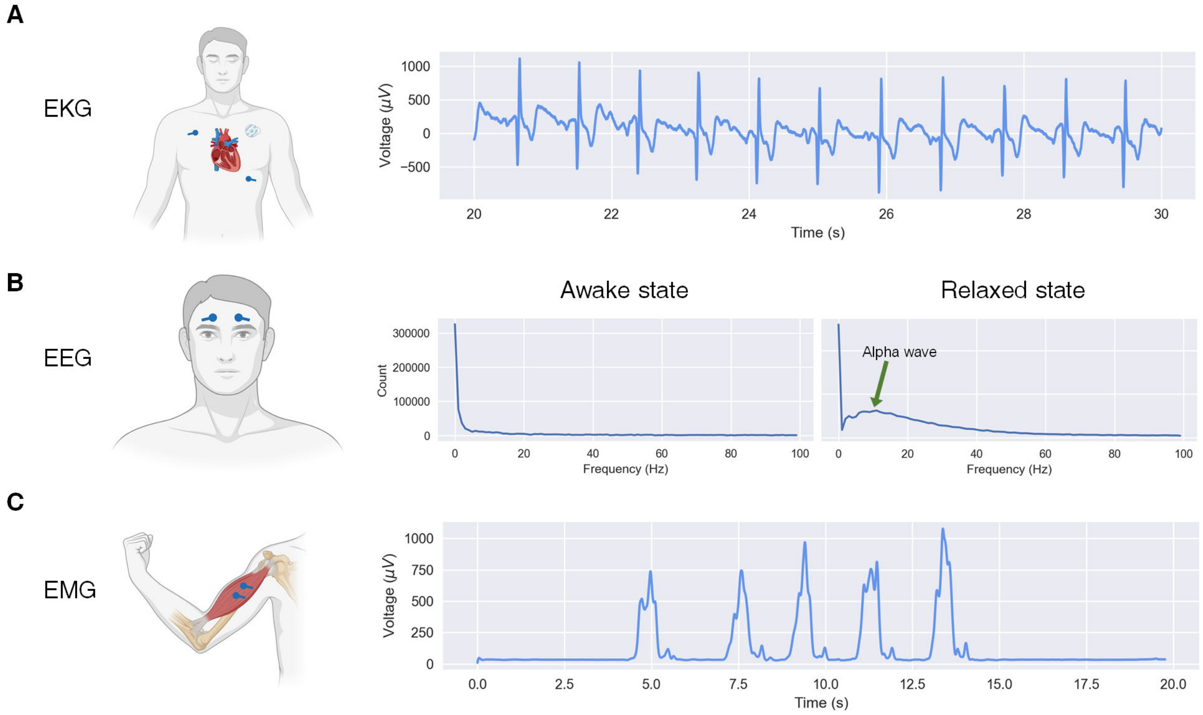


**Supplementary Figure S8** | Traces of electrocardiography (EKG), electroencephalography (EEG), and electromyography (EMG) signals detected using the Solution CP-G 250 electrodes, along with corresponding anatomical illustrations. The figures on the left depict the electrode placements for each modality. **A**, The left-side illustration shows the placement of three EKG electrodes on the chest in the conventional arrangement for monitoring heart electrical activity. The corresponding trace displays typical cardiac cycles with sharp R-peaks, confirming effective signal acquisition by the Solution CP-G 250 electrodes. **B**, The illustration represents the placement of three EEG electrodes on the forehead, aligned with the standard setup for detecting alpha waves, especially during relaxation exercises and controlled breathing. The EEG trace shows the frequency spectrum in both awake and relaxed states, with a clear appearance of alpha waves (8–12 Hz) during the relaxation phase, highlighting the precision of the Solution CP-G 250 in capturing subtle neurological signals. **C**, The illustration shows the placement of two EMG electrodes along the biceps muscle, used to measure electrical activity during muscle contractions. The corresponding trace shows muscle activation phases, with distinct peaks marking periods of biceps contraction, confirming the electrodes' ability to reliably detect muscle activity for applications such as myoelectric control or rehabilitation.

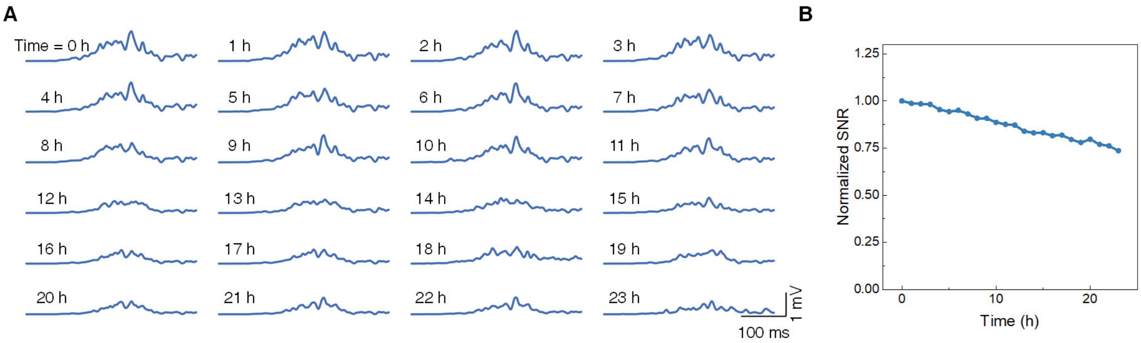


**Supplementary Figure S9 | Long-term sEMG recording stability over 24 hours.**
**A**. Representative sEMG recordings taken at 1-hour intervals over a 24-hour period, showing gradual signal degradation. **B**: Normalized SNR decrease, indicating a 25% reduction in amplitude after 24 hours of continuous use.

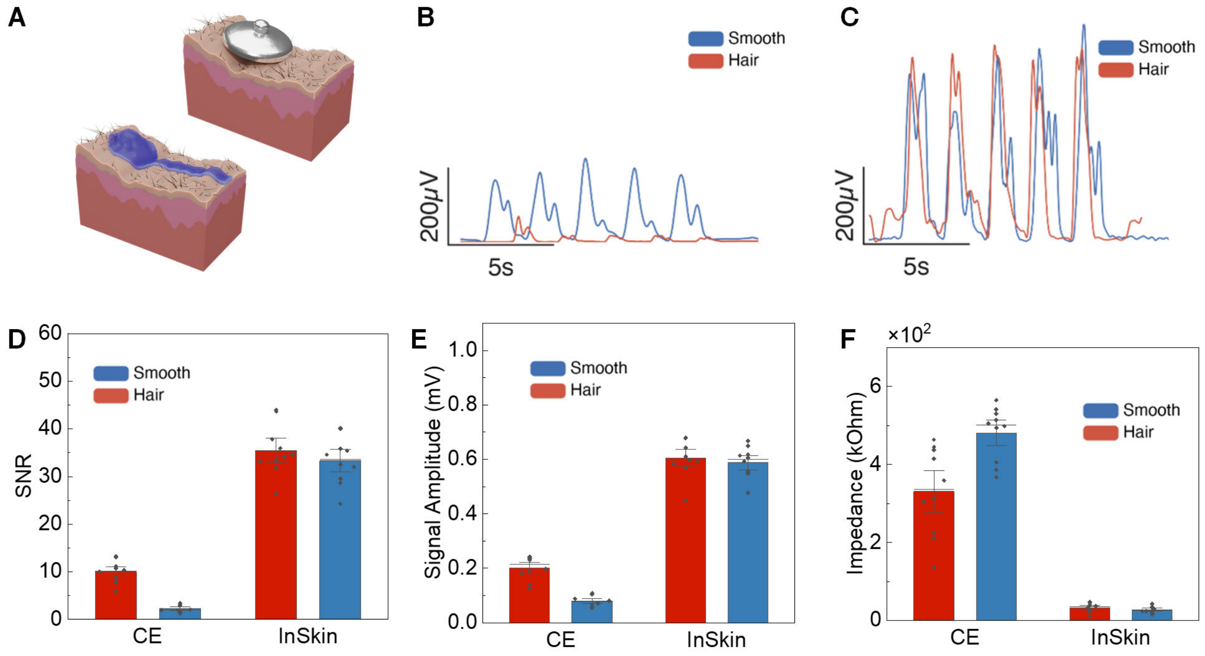


**Supplementary Figure S10 | Comparison of CE and CP-G 250 electrodes on smooth and hairy skin for sEMG recordings.**

**A.** Schematic illustration of the commercial electrode (CE) and solution CP-G 250 electrodes on hairy skin surfaces. **B, C**. Representative sEMG recordings from smooth (blue) and hairy (red) skin using CE (**B**) and CP-G 250 (**C**) electrodes. The CP-G 250 electrode demonstrates higher signal fidelity, particularly on hairy skin. **D.** SNR comparison between CE and CP-G 250 electrodes on smooth and hairy skin, showing significantly higher SNR for CP-G 250, particularly on hairy skin. **E**. Signal amplitude comparison, highlighting enhanced signal capture with CP-G 250 electrodes across both skin types. **F**. Electrode-skin impedance measurements, with CP-G 250 showing significantly lower impedance on both skin types, indicating superior skin-electrode interface stability.

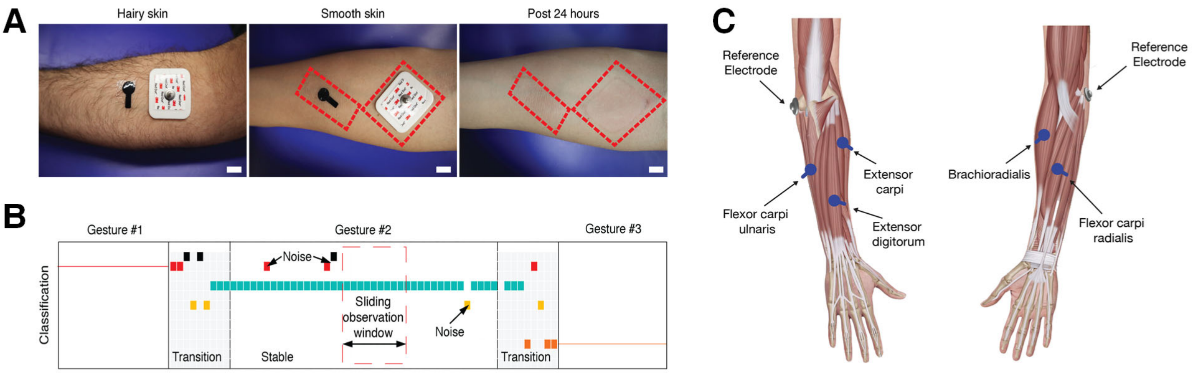


**Supplementary Figure S11** | **A,** Photos of commercially available EKG/EMG electrodes and Solution CP-G 250 on (1) hairy skin, (2) smooth and shaved skin, and (3) after their removal post-long-term comfort study. Unlike commercially available, the photos show how  Solution CP-G 250 electrodes can better conform to diverse skins and how they do not irritate the skin or leave residues upon sensor removal. **B**, Graphical representation of the custom algorithm employed for gesture recognition in sEMG prosthetic control. The EMG data for each channel is segmented into several nonoverlapping windows with widths of 1 second. When switching between gestures, the algorithm analyzes the signal in these 1-second windows to allow normalization. The window will be recognized and labeled as a valid gesture if the classification accuracy exceeds 0.7; otherwise, the data will be marked as a transition state and not recorded. A sliding observation window then compares the 1-second-long signal with specific criteria, and the predicted value is converted into finger angles. These angles are then sent to the prosthetic robotic hand system. **C,** Schematic of electrode placement around forearm muscles for optimized sEMG signal capture.
